# Allosteric inhibition of *Staphylococcus aureus* MenD by 1,4-dihydroxy naphthoic acid: A feedback inhibition mechanism of the menaquinone biosynthesis pathway

**DOI:** 10.1101/2022.07.12.499824

**Authors:** Tamsyn Stanborough, Ngoc Anh Thu Ho, Esther M. M. Bulloch, Ghader Bashiri, Stephanie S. Dawes, Etheline W. Akazong, James Titterington, Timothy M. Allison, Wanting Jiao, Jodie M. Johnston

## Abstract

Menaquinones (MKs) are electron carriers in bacterial respiratory chains. In *Staphylococcus aureus* (*Sau*), MKs are essential for aerobic and anaerobic respiration. As MKs are redox-active, their biosynthesis likely requires tight regulation to prevent disruption of cellular redox balance. We recently found that the *Mycobacterium tuberculosis* MenD, the first committed enzyme of the MK biosynthesis pathway, is allosterically inhibited by the downstream metabolite 1,4-dihydroxy-2-naphthoic acid (DHNA). To understand if this is a conserved mechanism in phylogenetically distant genera that also utilize MK, we investigated whether the *Sau*-MenD is allosterically inhibited by DHNA. Our results show that DHNA binds to and inhibits SEPHCHC synthase activity of *Sau*-MenD enzymes. We identified residues in the DHNA binding pocket that are important for catalysis (Arg98, Lys283, Lys309) and inhibition (Arg98, Lys283). Furthermore, we show that exogenous DHNA inhibits growth of *Sau*, an effect that can be rescued by supplementing the growth media with MK-4. Our results demonstrate that despite a lack of strict conservation of the DHNA-binding pocket between *Mtb*-MenD and *Sau*-MenD, feedback inhibition by DHNA is a conserved mechanism in *Sau*-MenD and hence the *Sau* MK biosynthesis pathway. These findings may have implications for the development of anti-staphylococcal agents targeting MK biosynthesis

## Introduction

Isoprenoid quinones such as menaquinones (MKs, vitamin K2) and ubiquinones (co-enzyme Q) are redox-active, membrane-bound molecules with vital roles across all domains of life (1). Menaquinones, considered to be the most ancient types of isoprenoid quinones, function as electron-carriers in bacterial electron transport chains, and are essential in ATP-generating redox reactions in mycobacteria, Gram-positive bacteria and anaerobically respiring Gram-negative bacteria. MKs also function as environmental sensors detecting changes in redox state and oxidative stress (2-8), and have been linked to biofilm formation in *Staphylococcus aureus* (*Sau*) (6), and latency and virulence in *Mycobacterium tuberculosis* (*Mtb*) (5, 9, 10).

MKs consist of a naphthoquinone head group linked to an isoprenoid tail of varying lengths and are therefore referred to as menaquinone-n (MK-n), where n denotes the number of isoprenyl side chain units. Side chain lengths of six to ten are common, but isoprenyl units of 1-5 and 11-15 are also found (2, 11). Two different pathways have evolved in bacteria for the biosynthesis of MK; the classical pathway (**Figure S1A**), utilised by bacteria such as *Escherichia coli, Mtb* and *Sau*, and the less common alternative (or futalosine) pathway (1, 2). Commonalities of both pathways include the separate synthesis of the naphthoquinone headgroup precursor and isoprenoid side chain. These components are then combined and further modified.

The MK biosynthesis pathway is absent in humans; and antibacterial drug discovery efforts targeting numerous enzymes from this pathway in bacterial pathogens have been reported (12-16). *Sau* is an important human pathogen, and utilises MK (predominantly MK-8) for both aerobic and anaerobic respiration (17-19). *Sau* is associated with a variety of skin, bone, joint and blood infections with considerable morbidity and mortality (20, 21). Treatment of *Sau* infections can be extremely challenging owing to the formation of persister populations and small colony variants (SCVs), high tolerance of *Sau* biofilm to antibiotics, and the emergence of antibiotic-resistant strains (22-25). These obstacles, in particular the appearance and spread of new drug-resistant strains, highlight the importance of identifying new inhibitors to treat *Sau* infections. Inhibitors of MenA, MenB and MenE enzymes in the *Sau* MK biosynthesis pathway exhibit promising growth inhibitory (26-30) and biofilm formation inhibitory activity (26), validating the druggability of this pathway for the development of *Sau*-targeting antibacterial agents.

Until recently, little was known about how the MK biosynthesis pathway is regulated. MK levels, and those of some upstream metabolites of the pathway, are likely to require tight regulation, as excessive concentrations of these molecules could result in cell toxicity if the redox balance is disrupted (2). We showed evidence of feedback regulation of the MK biosynthesis pathway in *Mtb* (31), by identifying the negative allosteric regulation of the first committed enzyme in MK biosynthesis, MenD by a downstream metabolite 1,4-dihydroxy-2-naphthoic acid (DHNA).

MenD is a thiamine diphosphate (ThDP)-dependent enzyme that catalyses the conversion of oxoglutarate and isochorismate to 2-succinyl-5-enolpyruvyl-6-hydroxy-3-cyclohexene-1-carboxylic acid (SEPHCHC) (**Figure S1**) (32, 33). Catalysis occurs through two covalent ThDP intermediates as the substrates oxoglutarate and isochorismate are successively added to ThDP before SEPHCHC is released (**Figure S1B**) (31, 34). The *Mtb-* MenD was found to bind DHNA in a pocket distinct from the active site in an arginine cage comprising Arg97, Arg277 and Arg303. All three arginine residues were identified as being crucial for full *Mtb*-MenD activity and for inhibition by DHNA (31). Although based on sequence similarity, the *Mtb*-MenD allosteric site was found to be well-conserved among mycobacteria and closely related *Rhodococcus sp*., the site was not strictly conserved in other microorganisms including *Sau*. This suggested that regulation of the MK biosynthesis pathway by DHNA may either be limited to a small group of bacteria, or that DHNA (or related molecules) may still bind in this region despite the absence of some key elements, such as the full arginine cage, found in the *Mtb*-MenD.

In this work, we found that the *Sau*-MenD is allosterically inhibited by DHNA. Thus, we demonstrate that inhibition by DHNA of the MK biosynthesis pathway is conserved between *Mtb* and *Sau*, which hints at an ancient origin of this allosteric feedback mechanism in Gram-positive bacteria. Furthermore, we show that exogenous DHNA inhibits growth of *Sau* bacteria, underscoring the importance of the MK pathway for growth. These findings suggest that small molecules based on the DHNA scaffold may be effective for the development of new anti-bacterial drugs.

## Material and methods

### Cloning, expression and purification of *Sau*-MenD

The MenD genes from *Sau* IS1 (sequence identical to gene locus tag EKM74_RS13930 in NCBI Reference Sequence NZ_AP019306.1) and *Sau* ATCC 25923 (gene locus tag KQ76_RS04840 in NCBI Reference Sequence NZ_CP009361.1) were amplified by PCR using MenD_Sau_F1 and MenD_Sau_R1 primers specified in **Table S1**. The amplified MenD genes were cloned into a modified pET30a vector using the restriction enzymes NcoI and HindIII to generate rTEV-cleavable N-terminally His_6_-tagged proteins. Cloning of alanine mutants (Arg98Ala, Lys283Ala and Lys309Ala) of the *S. aureus* IS1 MenD gene was performed by site-directed mutagenesis and In-fusion cloning using primers listed in **Table S1**.

His_6_-tagged *Sau*-MenD was expressed in *Escherichia coli* BL21 (DE3) cells. Cultures were grown for 18 h +/-2 h in Luria-Bertani Miller (LB) media (Thermo Scientific) with 50 μg/mL kanamycin at 37 °C. Seed cultures were diluted 1/200 in Terrific Broth autoinduction media (24 g/L yeast extract, 12 g/L tryptone, 0.8% v/v glycerol, 0.0162 M KH_2_PO_4_, 0.0528 M K_2_HPO_4_, 2 mM MgSO_4_, 0.375% w/v aspartic acid, 0.015% w/v glucose and 0.5% w/v lactose) containing 50 μg/mL kanamycin and cultured at 37 °C under agitation (180 rpm) for 3 h, followed by a further 20 h incubation at 18 °C and 180 rpm. Cells were harvested and lysed in lysis buffer (20 mM HEPES pH 8, pH 8, 150 mM NaCl, 5 mM CaCl_2_, 5% (v/v) glycerol, 50 mM imidazole and 1 mM TCEP) using a Microfluidics cell disrupter (Newton, MA). Recombinant *Sau*-MenD was purified at room temperature by IMAC with 5 mL HisTrap HP columns (GE Healthcare) and an imidazole gradient of 20–500 mM over 50 mL. Peak fractions were immediately diluted 1:2 in 1x lysis buffer to avoid protein precipitation. To remove the N-terminal His_6_-tag from the recombinant protein, TEV-cleavage (35) was performed overnight at 4 °C by dialyzing the protein in 1x lysis buffer without imidazole. Subtractive IMAC was then conducted at room temperature with 5 mL HisTrap HP columns to remove the cleaved His_6_-tag and rTEV. Fractions containing the target protein were combined and concentrated for further purification by size-exclusion chromatography (SEC). SEC was carried out at room temperature on a Superdex 200 10/30 column with buffer containing 20 mM HEPES pH 8, 150 mM NaCl, 5 mM CaCl_2_, 5% (v/v) glycerol and 1 mM TCEP. For long-term storage, 50% (v/v) glycerol was added to take the final concentration to 10% (v/v) and the protein solution was kept at -80 °C.

### Sequence alignment of MenD proteins

MAFFT sequence alignment of *Sau*-MenD_IS1_ and *Mtb*-MenD was performed in Geneious (36) using MAFFT version 7.388 (37), the AUTO algorithm, Blosum45 scoring matrix and a gap open penalty of 1.53 and an offset value of 0.153.

### Crystallisation

Initial *Sau*-MenD low resolution structures (incomplete and not presented in this work) were from both IS1 and ATCC wildtype (WT) enzymes in a range of MORPHEUS screen conditions (38) without and with ThDP present (1–2.5 mM). Optimised crystals for the higher resolution structure presented in this work were grown using tag-removed recombinant *Sau*-MenD_IS1_ (15 mg/mL in 20 mM HEPES pH 8, 150 mM NaCl, 5 mM CaCl_2_, 5% (v/v) glycerol, 2 mM TCEP and 2 mM ThDP) in 96-well sitting-drop format from MORPHEUS screen condition H12 (12.5% w/v PEG 1000, 12.5% w/v PEG 3350, 12.5% v/v MPD, 0.02 M of amino acids (sodium L and D-glutamate, L and D-alanine, glycine, L and D-lysine HCl, L and D-serine and 0.1 M bicine/Trizma base pH 8.5.). Crystals grew within 7 days and were flash frozen in liquid nitrogen.

### Data collection, structure determination and refinement

All diffraction data were collected using the macromolecular crystallography beamline MX2 at the Australian Synchrotron equipped with Dectris EIGER 16M detector (39). Data was autoprocessed via the Australian synchrotron pipeline, using iterative rounds of XDS auto-indexing (40, 41) and the resulting .hkl file was imported into the CCP4 program suite (42) for space group assignment, merging, truncating and generation of an R_*free*_ set of 5 % using *AIMLESS* (43). Analyses of merged CC½ correlations between intensity estimates from half data sets were used to influence high-resolution cutoff for data processing (44). Matthews co-efficient analysis (45) suggested 4 molecules per asymmetric unit (47.9 % solvent content).

The structures were solved by molecular replacement using Phaser (46), with the prior mentioned initial low resolution partially complete *Sau*-MenD model used as a search model (this model was generated from lower resolution *Sau*-MenD data using molecular replacement from *B. subtilis* (*Bs*)-MenD Protein Data Bank (PDB) code 2X7J (33) and manual model building/refinement).

The *Sau*-MenD IS1 model presented here was completed with iterative rounds of manual building using COOT (47) and refinement using Refmac5 (48) and Phenix (49). Additional density corresponding to ThDP, ions and glycine (from the crystallization medium) were modelled using available PDB dictionary restraints. Water molecules were identified by their spherical electron density and appropriate hydrogen bond geometry with the surrounding structure. The final refined 2.35 Å structure was deposited in the PDB with the code 7TIN. Unless otherwise stated, all protein structure images were generated using PyMOL (The PyMOL Molecular Graphics System, Version 1.5, Schrödinger, LLC).

### Native mass spectrometry

Protein samples for native mass spectrometry (*Sau*-MenD_IS1_ 4.7 mg/mL, and *Sau*-MenD_ATCC 25923_ 6.3 mg/mL) were exchanged from size exclusion buffer into 2 M ammonium acetate pH 7.4 via a single-pass through a Bio-Spin P-6 column (Bio-Rad). Buffer-exchanged protein was loaded into gold-coated glass capillaries (1.0 mm OD/0.75 mm inner diameter with filament), fabricated with orifice sizes of ≈5 μm, using a P-2000 puller (Sutter). Spectra were acquired using a Synapt XS (Waters) equipped with a 32k quad, with instrument settings optimised for the transmission of high-mass ions, and soft ionisation and activation conditions (50, 51). The instrument was mass calibrated using CsI. Spectra were smoothed (3×20, mean) and mass assigned in MassLynx (Waters).

### Intrinsic fluorescence quenching experiments

Intrinsic fluorescence quenching experiments were performed at 25 °C with a Varian Cary Eclipse Fluorescence Spectrophotometer (Agilent Technologies) with slits set at 5 nm bandwidth. The excitation wavelength was 290 nm and emission spectra were recorded in the 300–400 nm range. Experiments were conducted in 3 mL quartz cuvettes with 1 μM *Sau*-MenD in an assay buffer consisting of 50 mM HEPES pH 8, 150 mM NaCl, 5 mM CaCl_2_ and 1 mM TCEP. For *Sau*-MenD samples in the presence of the reaction ligands, the enzyme was preincubated with 100 μM ThDP only or 100 μM ThDP and 300 μM oxoglutarate for 45 min at ambient temperature. DHNA (Sigma-Aldrich) was titrated from stock solutions that were prepared in the assay buffer and kept on ice. Following DHNA addition, cuvette contents were mixed immediately and fluorescence intensity was measured.

The fluorescence intensities obtained at 340 nm were corrected for the dilution factor as well as for inner filter effects (52) using equation 1,

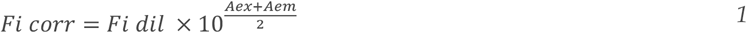

where *Fi corr* is the corrected value of fluorescence intensity at the titration point, *Fi dil* is the dilution-corrected fluorescence intensity measured, and *Aex* and *Aem* are the absorbance of DHNA at excitation (290 nm) and emission wavelengths (340 nm).

Corrected intensities were plotted against the DHNA concentration and to determine the K_d_ value for DHNA, the resulting data was fitted in GraphPad Prism 9 using equation 2,

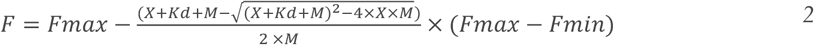

where *F* is the measured fluorescence intensity of *Sau*-MenD, *Fmax* is the fluorescence intensity of *Sau*-MenD in the absence of DHNA, *Fmin* is the fluorescence intensity when *Sau*-MenD is saturated with DHNA, *M* is the μM concentration of enzyme used in the assay and *Kd* is the dissociation constant.

### UV spectroscopy assays

The isochorismate mixture was prepared using commercially produced chorismic acid and laboratory made *E. coli* MenF as described in Bashiri et al. 2020 (31, 32). *Sau*-MenD activity and DHNA-inhibition assays were performed using the UV spectroscopy assay described by Bashiri et al., 2020 (31) with some modifications. The assay buffer consisted of 50 mM HEPES pH 8, 150 mM NaCl and 5 mM CaCl_2_. Reaction mixes consisting of 0.4 μM *Sau*-MenD, 100 μM ThDP and 300 μM oxoglutarate were incubated in the reaction buffer at 37 °C for 30 min prior to adding 20 μM isochorismate to initiate the reaction. All assays were carried out using a Cary 400 UV-Vis spectrophotometer (Agilent Technologies) and quartz cuvettes with a final reaction volume of 150 μL. Initial rate data were fitted using the Cary WinUV software.

DHNA-inhibition assays contained 0.4 μM protein, 100 μM ThDP, 300 μM oxoglutarate and various concentrations of DHNA (0–51.2 μM). After pre-incubation of these components at 37 °C for 30 min, 10 μM isochorismate was added to initiate the reactions. Stock solutions of DHNA were freshly prepared in assay buffer prior to each experiment and stored on ice. At 25.6 and 51.2 μM DHNA, an increase in absorbance was observed for control samples run without isochorismate owing to gradual DHNA oxidation. To account for this, the linear increase in absorbance at 278 nm of the non-isochorismate controls was deducted from the initial rate data of these samples. Initial rate data was then fitted to the four-parameter logistic Hill equation with GraphPad Prism 9.

### Growth assays with DHNA and MK-4 rescue assays

*Sau* and *Pseudomonas aeruginosa* (**Table S2**) were grown for 18 h +/-2 h in 5 mL of LB media at 37 °C and 180 rpm. Cultures were then diluted in fresh LB media to contain 2.7 × 10^6^ CFU/mL. A Nunc Microwell 96-well flat-bottom plate (Thermo Scientific) was seeded in triplicate (from three independent starter cultures) with 75 μL of each of the diluted cultures and 75 μL of 2x working solutions of DHNA, MK-4 or DHNA + MK-4 (prepared from the following stock solutions: 100 mM DHNA dissolved in DMSO and 50 mM MK-4 [Sigma-Aldrich] dissolved in 100% ethanol). Plate wells contained a final concentration of 10^6^ CFU/well with 0, 50 or 100 μM DHNA for growth assays; or with 0 or 100 μM DHNA supplemented with 337 μM MK-4 for growth rescue assays. Plates were incubated at 37 °C and 300 rpm for 24 h. Growth was determined by measuring the OD600 of the cultures every 30 min over the incubation period. Data from biological replicates was averaged and subtracted from blank data (sterile media with appropriate additives).

## Results

The *Sau*-MenD shares 28% identity with its well-studied homologue in *Mtb* (*Mtb*-MenD) (**Figure S2**) and eluted in one peak in SEC at an elution volume near identical to the *Mtb*-MenD protein (data not shown). The tetrameric conformation of this protein was confirmed by native mass spectrometry (**Figure S3**).

### *Sau*-MenD binds DHNA

To determine whether the *Sau*-MenD can bind DHNA, the intrinsic fluorescence of MenD enzymes from *Sau* IS1 and *Sau* ATCC 25923 was monitored upon DHNA titration in the presence and absence of the co-factor ThDP, and in the presence of ThDP and first substrate oxoglutarate. The two *Sau*-MenD variants differed in sequence by seven amino acids and were therefore used to assess potential differences in their ability to bind DHNA. Although there are no tryptophan residues present in the DHNA binding sites of either *Sau*-MenD variant (based on the *Mtb*-MenD DHNA binding site), each of the enzymes contains four tryptophans, one of which is near the pocket (Trp293).

An emission maximum at 340 nm resulted from excitation of the *Sau*-MenD proteins at 290 nm. Fluorescence quenching upon DHNA titration had saturation behaviour and was fitted to equation 2 to estimate the *K*_d_ values (**Figure 1**). Our results show that the apoenzyme, ThDP-bound and ThDP- and oxoglutarate-bound forms of *Sau*-MenD bind DHNA with *K*_d_ values between 58–110 μM for the IS1 protein and 67–85 μM for the ATCC 25923 variant. The affinity of the ThDP- and oxoglutarate-bound form of *Sau*-MenD_IS1_ for DHNA (*K*_d_ = 58 μM, 95% CI 38–87 μM) is slightly higher than that of the apoenzyme (*K*_d_ = 110 μM, 95% CI 91–150 μM). While the *K*_d_-values for the apoenzyme (*K*_d_ = 85 μM) versus the ThDP- and oxoglutarate-bound states (*K*_d_ = 67 μM) of *Sau*-MenD_ATCC 25923_ show a similar trend, overlapping 95% confidence intervals hinder a definitive conclusion.

**Figure 1.**
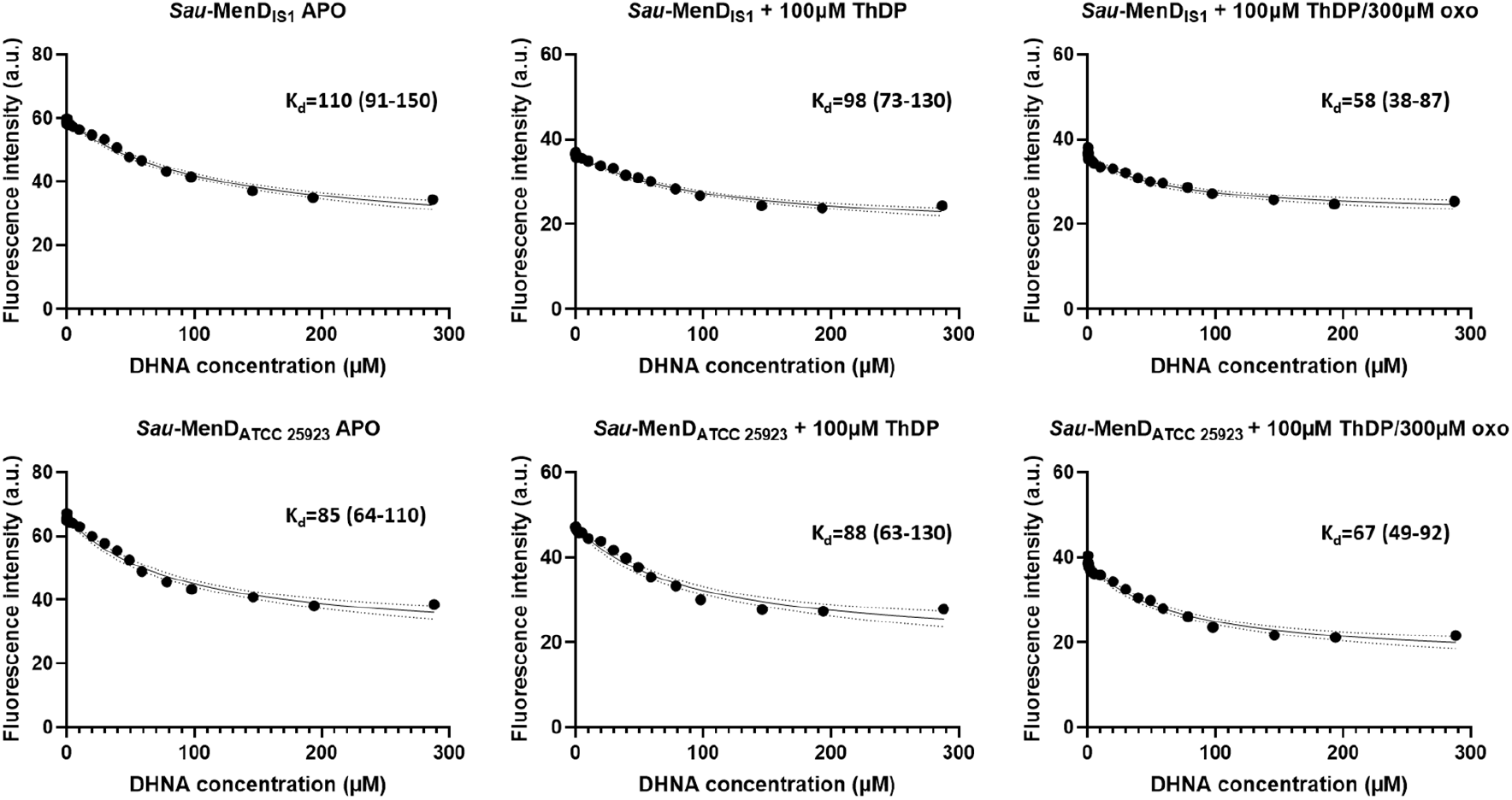
Intrinsic fluorescence quenching of *Sau*-MenD enzymes upon DHNA titration. Intrinsic fluorescence quenching experiments were performed with 1 μM enzyme. For *Sau*-MenD samples complexed with 100 μM ThDP only, or 100 μM ThDP and 300 μM oxoglutarate (oxo), enzymes were preincubated with the respective ligands for 45 min at ambient temperature. DHNA was then titrated, cuvette contents were mixed immediately and fluorescence intensity was measured twice for each titration. The fluorescence intensities obtained at 340 nm were corrected for the dilution factor and inner filter effects. To obtain dissociation constants for DHNA, the resulting data were fitted in GraphPad Prism with equation 2. *K*_d_ values are shown in μM and respective 95% confidence intervals are provided in parentheses.

### DHNA inhibits *Sau*-MenD SEPHCHC synthase activity

We previously showed that DHNA inhibits the SEPHCHC activity of 0.6 μM *Mtb*-MenD with an IC_50_ of 53 nM (31). To determine whether this inhibitory mechanism is conserved in *Sau*-MenD, the effect of DHNA on the MenD enzymes from *Sau* IS1 and *Sau* ATCC 25923 was investigated in a UV spectroscopy-based assay by measuring isochorismate consumption at 278 nm in the presence of various concentrations of DHNA (0–51.2 μM). Our assay results (**Figure 2 and Table 1**) show that DHNA inhibited 0.4 μM *Sau*-MenD_IS1_ and *Sau*-MenD_ATCC 25923_ with IC_50_ values of 3.7 μM and 2.3 μM, respectively, suggesting this last soluble metabolite of the pathway is also an allosteric regulator of the *Sau*-MenD.

**Figure 2.**
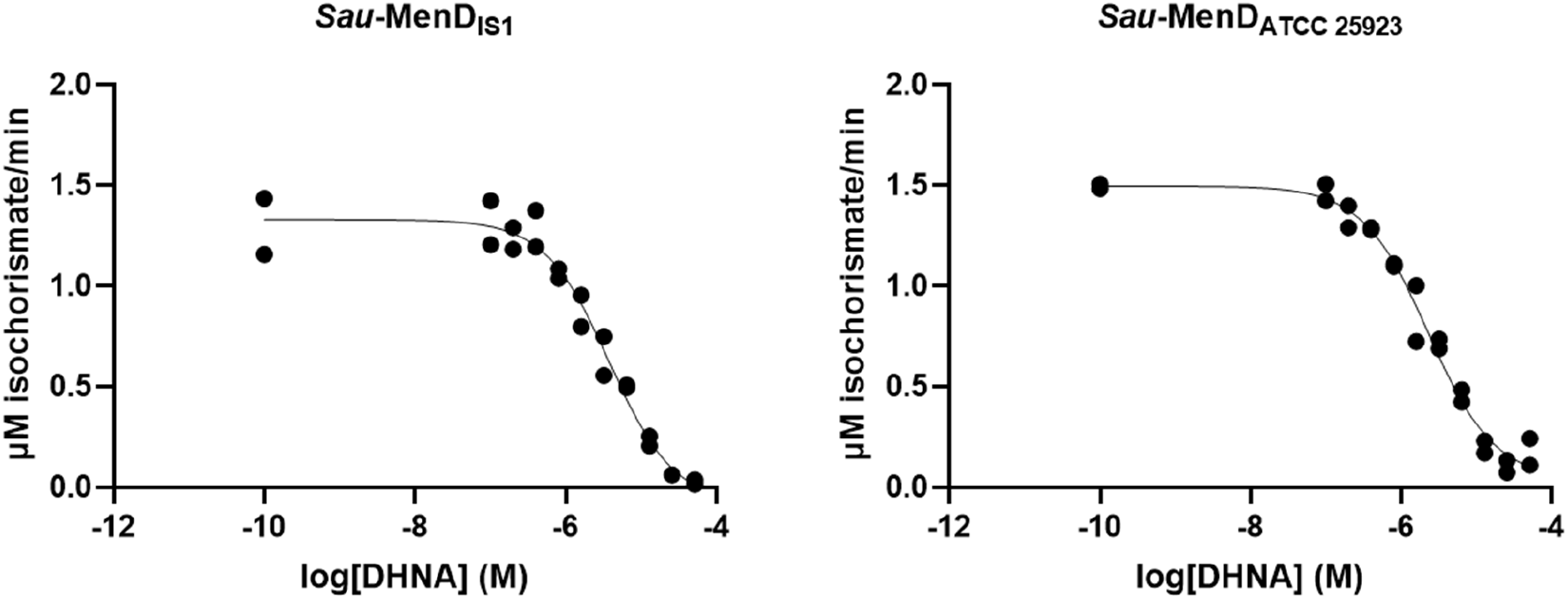
Inhibition of *Sau*-MenD enzymes by DHNA. Using a UV spectroscopy-based assay for isochorismate consumption, inhibition assays were performed with 0.4 μM enzyme, 100 μM ThDP, 300 μM oxoglutarate and various concentrations of DHNA (0.1–51.2 μM). Following a 30 min preincubation of enzyme with ThDP, oxoglutarate and DHNA, 10 μM isochorismate was added to initiate the reaction. Initial rates were measured and fit to the four-parameter logistic Hill equation. Data shown for each enzyme are from two independent experiments.

**Table 1.**
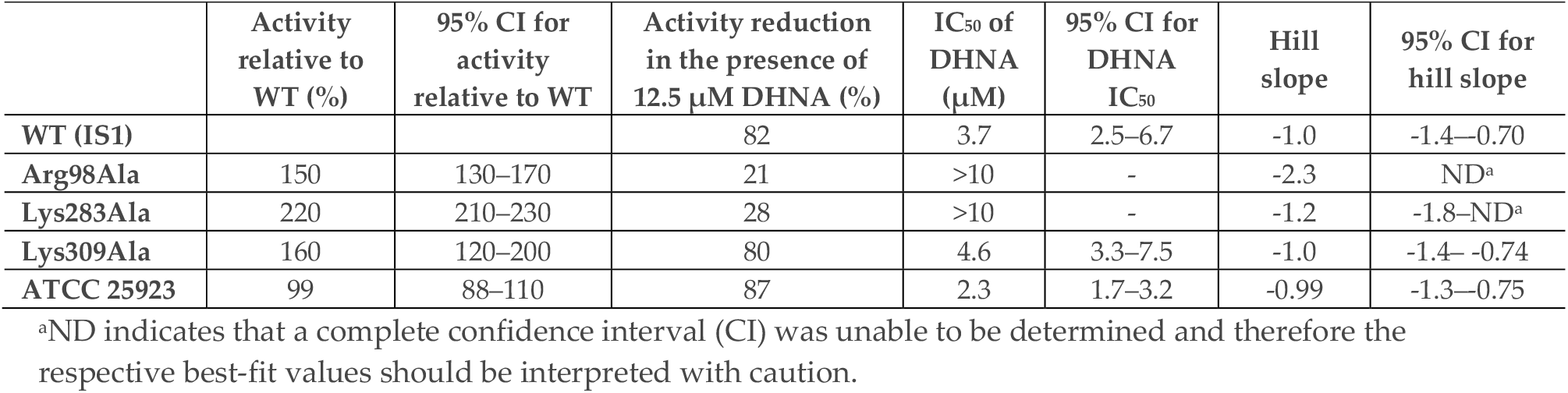
Activity and DHNA inhibition of *sau*-MenD variants.

### Structure of *Sau*-MenD

The *Sau*-MenD_IS1_ structure was solved (*processing and refinement statistics* **-Table S3**) to a resolution of 2.35 Å in space group *P*2_1_2_1_2_1_ with four chains (A–D, with interpretable density across their length for residues 2– 554/557) in the asymmetric unit comprising the biological “dimer of dimers” tetrameric unit (**Figure 3A**), consistent with our native MS analysis (**Figure S3**). The *Sau*-MenD_IS1_ tetramer structure is typical of all known MenD structures to date and, like the *B. subtilis* (*Bs*)-MenD (33) and *E*.*coli* (*Ec*)-MenD (53) structures, is symmetrical with nearly identical conformations for each monomer (RMSD using SSM (54) of 0.29–0.44 Å over the length of the chain) and well-defined ThDP and Ca^2+^ cofactors bound to all four active sites.

**Figure 3.**
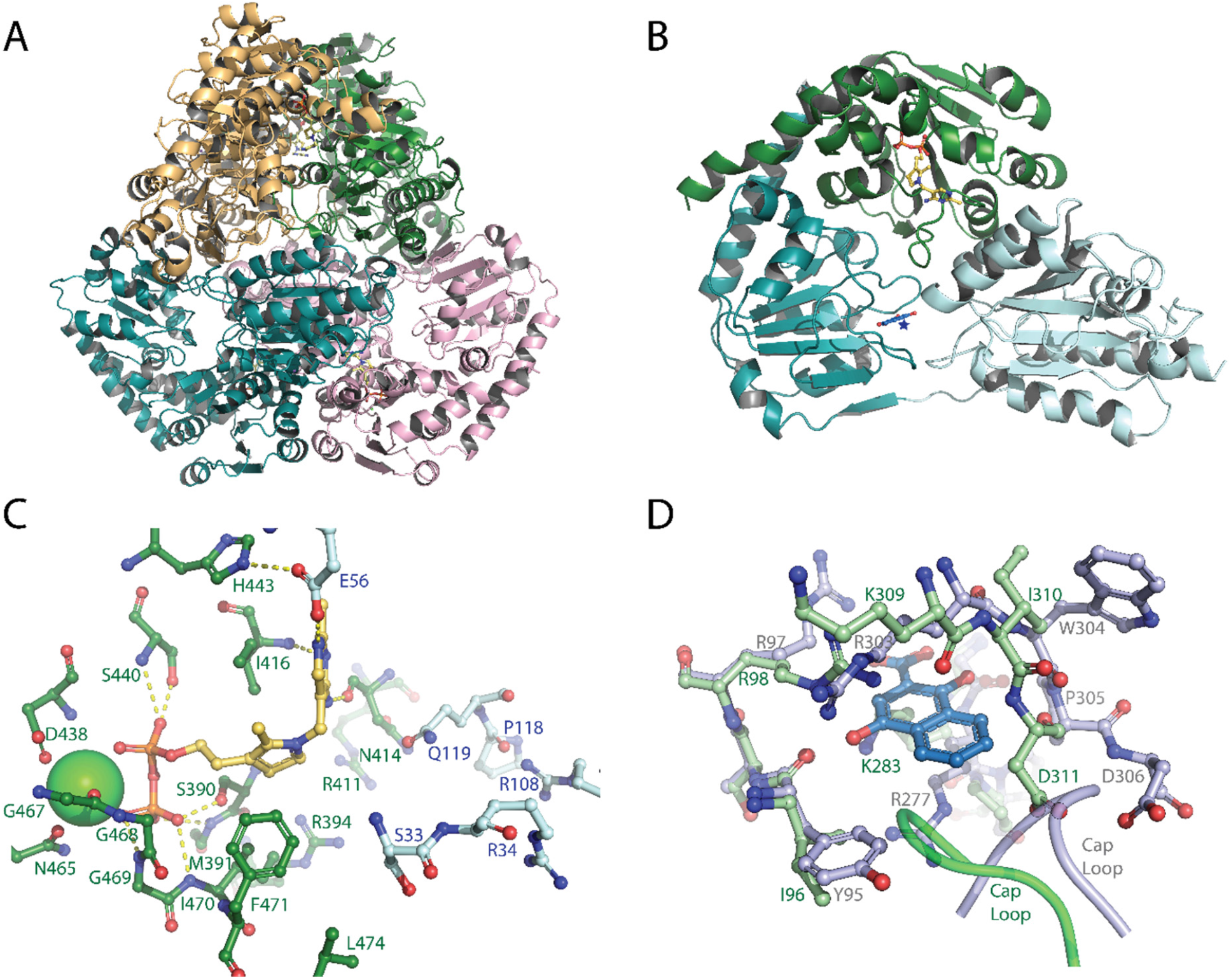
Structure of *Sau*-MenD. **A**, Cartoon depiction of the tetrameric “dimer of dimers” present in the asymmetric unit. Dimer 1 comprises the green and orange chains, dimer 2 the teal and pink chains and the four ThDP in the active sites are shown as yellow sticks. **B**, Cartoon depiction of the three-domain monomer fold. Domain I [PYR] is shown in light blue, domain II [TH3] in teal, domain III [PP] in green and ThDP is shown as yellow sticks with the associated Ca^2+^ ion as a green sphere. The location of the DHNA binding site from *Mtb*-MenD is depicted by a star and the *Mtb*-MenD DHNA from the overlay in D is shown in blue sticks. **C**, A close-up of the *Sau*-MenD active site. ThDP is shown as yellow sticks with the associated Ca^2+^ ion as a green sphere. Active site residues are shown as sticks (domain III (PP) is green with green labels and domain I (PYR) is light blue with blue labels) and selected hydrogen bonds are shown as yellow dashes. **D**, Overlay of the *Mtb*-MenD DHNA binding site (grey-blue sticks with grey labels) with DHNA bound (blue sticks) and the *Sau*-MenD putative allosteric site (green sticks with green labels).

The three-domain α/β monomeric fold (**Figure 3B**) is typical of enzymes from the decarboxylase superfamily of ThDP-dependent enzymes (55), with domain I (PYR domain, residues 1–200) pairing with domain III (PP domain, residues 364–557) from the other monomer in the dimer to form each of the four paired MenD active sites. Similar to *Bs*-MenD (33), the extended *Sau*-MenD_IS1_ C-terminal helix at the end of domain III extends down, packing against a domain III active site lid (residues 469–489) before ending in domain II (TH3, residues ∼ 220–364) (**Figure 3B**). Domain II (TH3), the most sequence-diverse of the MenD domains, is connected to domain I via a long linker (∼ residues 201–220) and to domain III via a long helix (residues 341-364). Domain II, which has no known function in many decarboxylase superfamily enzymes, has been associated with formation of the allosteric regulatory DHNA binding site in *Mtb*-MenD (31). Despite the overall modest sequence conservation across MenD enzymes, the *Sau*-MenD_IS1_ active site shows high conservation with other well-studied MenD enzymes for the residues involved in ThDP, substrate/intermediate binding and catalysis (**Figure 3C**). The ThDP diphosphate and associated Ca^2+^ are anchored to the enzyme via domain III (Ser390, Met391; Ser440, Asp438, Gly’s 467-469, Asn465, Ile471), as is the ThDP thiazole ring which is sandwiched between the side chains of Phe471 and Ile416, facilitating the catalytically active V-shaped ThDP conformation (**Figure 3C**). The ThDP aminopyrimidine ring inserts into a pocket between domain I and domain III forming hydrogen bonds with Asn414, Ile416 and Glu56; the latter playing an important role in proton transfer and stabilization of the various catalytically important aminopyrimidine ring tautomers (32, 34).

Substrate/intermediate binding residues come from both domain I and III; intermediate I (Arg394, Arg411) and intermediate II (Ile470, Phe471 and Leu474) interacting residues from domain III; carboxylate/decarboxylation binding pocket residues Ser33, Arg34 and intermediate II interacting residues Arg108 and Glu119 from domain I (**Figure 3C**) (31, 32). Invariant in MenD enzymes, Glu119 has been ascribed several important roles during catalysis, including product release (31, 32). Both Arg108 and Glu119 reside on a mobile loop shown in *Mtb*-MenD to connect the active site and allosteric sites (loop residues 114–116 cap the allosteric site (**Figure 3D**)) (31). This loop normally contains another key intermediate I/II binding residue (equivalent to *Mtb*-MenD Asn117) but in *Sau*-MenD this is replaced by a proline, and the side chain of Asn414 from domain III takes on a structurally similar role, similar to *Bs*-MenD (32, 33).

In contrast to the high conservation at the active site, conservation of the *Mtb*-MenD DHNA-binding allosteric site across MenD enzymes appears limited. Sequence comparisons with *Sau*-MenD indicated low conservation of the allosteric region, with the potential for conservation of one of the arginine cage residues and substitution of another for a Lys (*Sau*-MenD Arg98 and Lys283). Attempts to co-crystallise DHNA with *Sau*-MenD resulted in precipitate, while soaking greatly impaired diffraction. Therefore, structural comparisons of the *Sau*-MenD structure and DHNA-bound *Mtb*-MenD were used to gain insight into the structural conservation at the putative allosteric site. These overlays reveal that the likely structural equivalents for the arginine cage are Arg98, Lys283 and Lys309 (**Figure 3D**). None of these residues match the sidechain conformers that would be needed for DHNA binding, suggesting that for DHNA to bind, conformational rearrangements would be needed along with backbone movements around the 305–312 region. In the *Sau*-MenD structures, Arg98 is extensively hydrogen bonded to the backbone of Asn308, Lys309 and the Asn308 sidechain. Interaction and stacking with DHNA would interfere with these interactions and could facilitate movement of the region to accommodate DHNA.

### Residues in the DHNA binding pocket are important for activity and inhibition

To validate our structural findings and investigate whether the equivalent arginine-cage candidate residues in *Sau*-MenD (Arg98, Lys283 and Lys309) are important for activity and DHNA inhibition, we performed alanine mutagenesis experiments of the WT *Sau*-MenD_IS1_. Interestingly, all three mutants showed enhanced activity compared to the WT *Sau*-MenD_IS1_ when measured under the same conditions and in the absence of DHNA (**Figure 4A, Table 1**). Mean activity compared to the WT was 150%, 220%, and 160% for the Arg98Ala, Lys283Ala, and Lys309Ala mutants, respectively, suggesting all three residues play an important role in catalytic activity, as was shown for the equivalent residues in *Mtb*-MenD.

**Figure 4.**
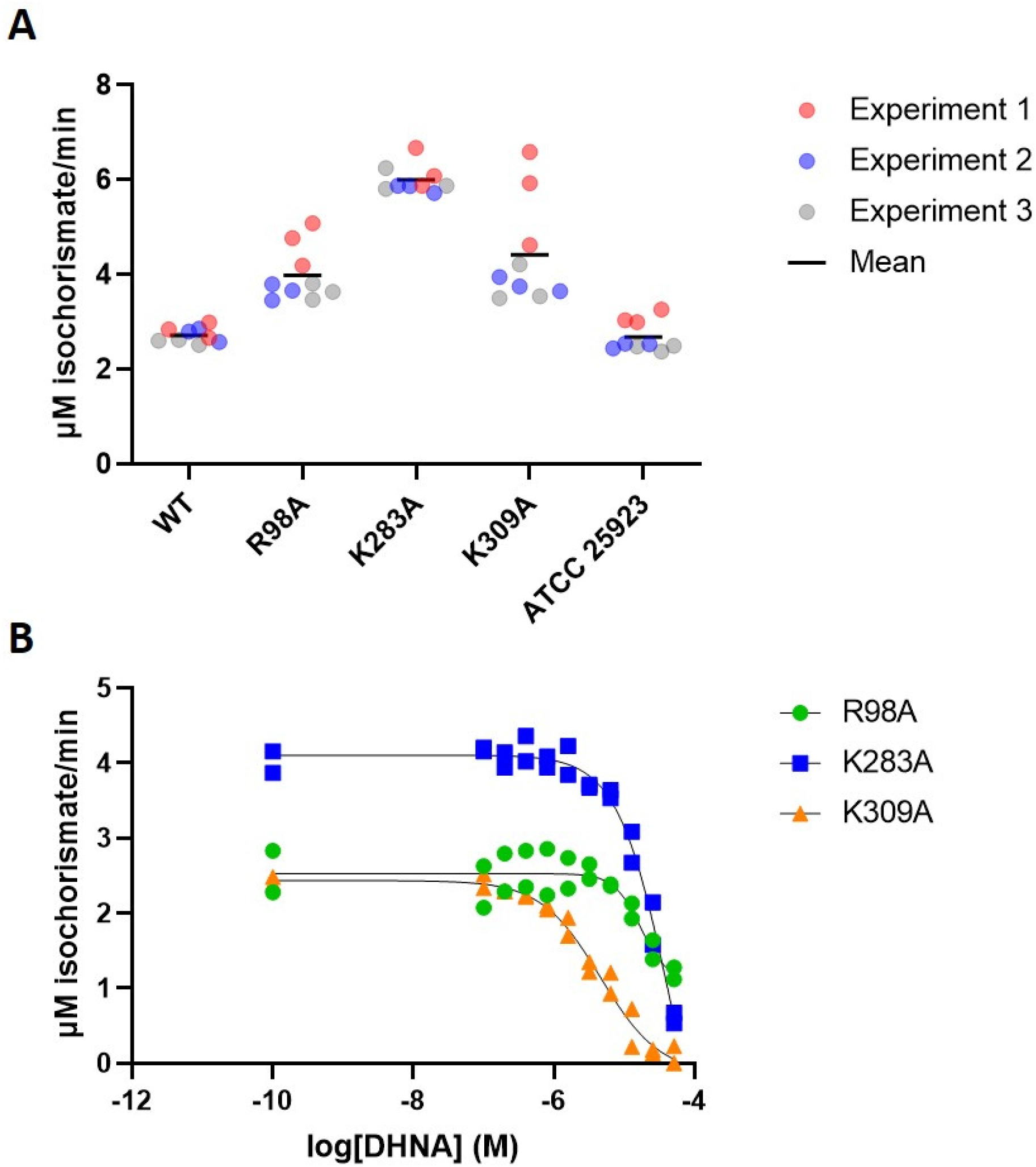
Activity and DHNA inhibition of *Sau*-MenD mutants. Using a UV spectroscopy-based assay for isochorismate consumption, activity (in the absence of DHNA) and DHNA inhibition of *Sau*-MenD alanine mutants was investigated. **A**, Activity of *Sau*-MenD mutants compared to WT *Sau*-MenD_IS1_ and *Sau*-MenD_ATCC 25923_. Three independent experiments were performed in technical triplicate and initial rate data were determined. **B**, IC_50_ data for DHNA against *Sau*-MenD mutants. Data shown for each enzyme are from two independent experiments. Initial rates were measured and fit to the four-parameter logistic Hill equation. Assays were performed with 0.4 μM enzyme, 100 μM ThDP and 300 μM oxoglutarate. DHNA (0.1–51.2 μM) was present for inhibition assays only. Following a 30 min preincubation of enzyme with ThDP, oxoglutarate and DHNA (inhibition assays only), 20 μM (or 10 μM for inhibition assays) isochorismate were added to initiate the reaction.

IC_50_ experiments revealed that DHNA does not inhibit the Arg98Ala and Lys283Ala mutants, as effectively as it does the WT protein (**Figure 2 and 4B, and Table 1**). DHNA has strong absorbance at 278 nm, thus it was not possible to test DHNA concentrations above 51.2 μM. This led to an inability to accurately determine IC_50_ values >10 μM and truncated inhibition curves for the Arg98Ala and Lys283Ala mutants (**Figure 4B**). As such, we were unable to determine IC_50_ values for these mutants. The truncated inhibition curves of the Arg98Ala and Lys283Ala mutants are an indication that these residues play an important role in DHNA inhibition. By contrast, the Lys309Ala mutant remained sensitive to DHNA inhibition with an IC_50_ of 4.6 μM, similar to that of the WT.

To illustrate the differences in DHNA-sensitivity between the mutant and WT enzymes, the percentage inhibition in activity at 12.5 μM DHNA was also determined (**Table 1**). This was the concentration at which the greatest differences in sensitivity were observed between the WT and mutant enzymes and it was the highest concentration of DHNA tested that did not require correction of the initial rate data owing to gradual DHNA oxidation. At 12.5 μM DHNA, the catalytic activities of the WT enzyme and Lys309Ala mutant were reduced by 82% and 80%, respectively, whereas the activities of the Arg98Ala and Lys283Ala mutants were notably less affected with reductions of 21% and 28%, respectively.

### DHNA supplementation inhibits growth of *Sau* in media

As the first metabolite in the MK biosynthesis pathway with a complete naphthoquinol ring, DHNA levels in the cells are likely to require tight regulation, as excessive amounts could result in cell toxicity if the redox balance is disrupted. To test this premise, four different *Sau* strains, including methicillin-resistant and methicillin-sensitive clinical isolates, were grown in media supplemented with various concentrations of DHNA (0–150 μM). *Pseudomonas aeruginosa* was used as a control as this bacterium does not produce menaquinone, only ubiquinone, and therefore does not contain classical MK biosynthesis genes (56). We found that exogenous DHNA at 50 μM clearly impaired growth of all four *Sau* strains (**Figure 5A–D**). The growth impact was more severe at 100 μM DHNA, and at 150 μM DHNA, growth of all four *Sau* was completely inhibited for the duration of the assay (24 h). By contrast, growth of *P. aeruginosa* was unaffected at DHNA concentrations up to 150 μM (**Figure 5E**).

**Figure 5.**
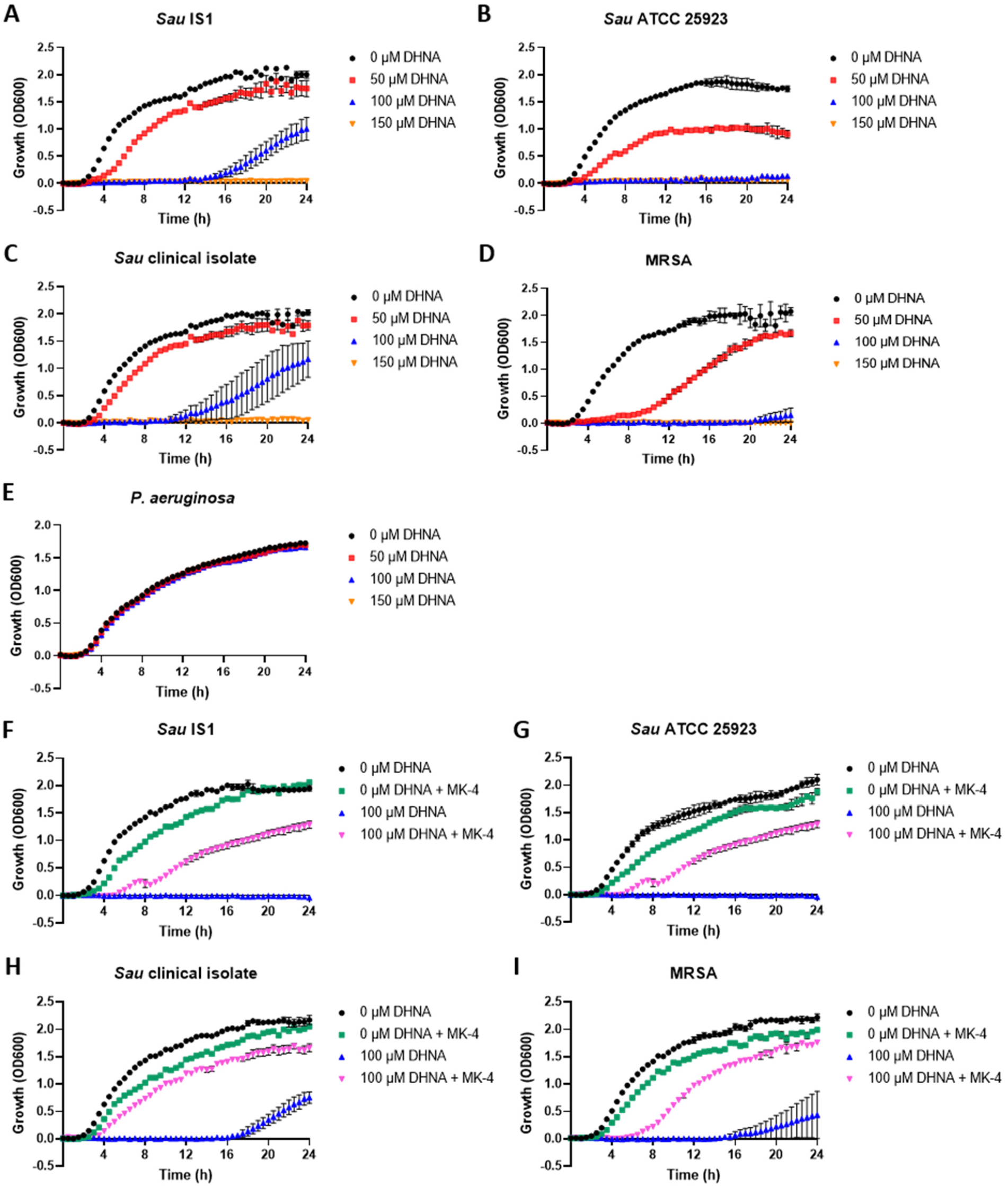
Growth assays with DHNA and MK-4 rescue assays. A–E Exogenous DHNA inhibits growth of *Sau*. Bacteria were grown at 37 °C under agitation in LB media containing various concentrations of DHNA (0, 50, 100 and 150 μM). **F–I MK-4 rescues growth of DHNA-treated *Sau***. *Sau* strains were grown at 37 °C under agitation in LB media with (100 μM) and without DHNA and supplemented with 337 μM MK-4. Growth for both assays was monitored over 24 h by OD600 measurements taken every 30 min. Data represent the mean ± SD of three biological replicates.

### MK-4 rescues growth defects caused by exogenous DHNA

To ascertain whether growth inhibition caused by DHNA is due to inhibition of the MK biosynthetic pathway, rescue experiments were performed by adding 337 μM MK-4 (150 μg/mL MK-4) to culture media with (100 μM DHNA) and without DHNA present, and monitoring growth of the *Sau* bacteria over 24 h. In the absence of DHNA, addition of MK-4 slightly impacted growth of the bacteria (**Figure 5F–I**). This may be due to oxidative stress, as MK is a redox-active metabolite. Despite this, MK-4 addition to culture media containing 100 μM DHNA resulted in a significant growth improvement of the strains compared to growth in the presence of 100 μM DHNA only. Notably, growth was not completely restored to levels observed in the absence of DHNA, which is possibly because MK-4 is not the preferred MK for *Sau* (predominant MK in *Sau* is MK-8).

## Discussion

We recently discovered feedback regulation of the MK biosynthesis pathway, with the last cytosolic metabolite of the pathway, DHNA, found to allosterically inhibit *Mtb-*MenD (31). Sequence alignments of MenD homologues suggested there was limited conservation of this allosteric site in other bacteria including the pathogen, *Sau*. However, data presented here demonstrate that DHNA is also able to bind to and inhibit the SEPHCHC synthase activity of *Sau*-MenD enzymes.

To better understand the basis for the conservation of allosteric inhibition of *Sau*-MenD by DHNA we solved the *Sau*-MenD structure and investigated conservation at a structural level. Like the *Mtb*-MenD, the *Sau*-MenD structure revealed a tetrameric protein, consisting of a dimer of dimers with paired active sites. Consistent with the full occupancy of the symmetrical *Ec* and *Bs*-MenD structures, and in contrast to the half-sites occupancy of the asymmetrical structures of *Mtb*-MenD, we did not see asymmetry in the *Sau*-MenD co-factor bound structure, and all four active sites were occupied with ThDP. There is limited conservation of the allosteric region; we identified only one of three key arginine’s of the *Mtb*-MenD arginine cage (Arg97) conserved in the *Sau*-MenD (Arg98) with the other two (*Mtb*-MenD Arg277 and Arg303) being conservatively replaced by Lys283 and Lys309.

These differences in the *Sau*-MenD allosteric site cage were reflected in notably higher IC_50_ values for DHNA in the low micromolar range, compared to low nanomolar range for the *Mtb*-MenD (31). Mutation of the three *Sau*-MenD allosteric cage candidates, validated a significant role for two (Arg98 and Lys283) in inhibition of SEPHCHC synthase activity by DHNA. The inhibitory potency of DHNA remained relatively unaffected by mutation of Lys309, suggesting this residue plays a much more subtle role in DHNA binding and inhibition.

In *Mtb*-MenD, Arg97 (the residue with hydrogen bond and electrostatic interactions with the DHNA carboxylate) is the arginine cage residue whose mutation results in the most impacted change in DHNA response. It is also the most conserved residue in sequence alignments (31). Our results validate the importance of this arginine (Arg98 in *Sau*-MenD) for allosteric regulation. Furthermore, they show that regulation, albeit at different potencies, can be conserved across bacterial species despite significant sequence divergence. This may reflect divergent needs of the bacteria. In contrast to *Mtb*, which relies on oxidative phosphorylation for growth (57, 58), substrate-level phosphorylation of fermentable sources allows *Sau* to grow slowly. In fact, respiration-deficient *Sau*, known as SCVs, have a slow-growth phenotype as a result of resorting to energy generation via fermentation. Therefore, *Mtb* and *Sau* may have different needs for regulation of this pathway, requiring adaptations of the allosteric site.

Assuming DHNA adopts a similar binding pose in *Sau*-MenD, our overlays with the *Mtb*-MenD structure suggest that several residues in the *Sau*-MenD allosteric site (all three cage residues and residues 305–314) would need to undergo movement to accommodate DHNA. It is plausible that initiation of DHNA binding facilitates a series of structural changes that help to achieve this. In the *Sau*-MenD structure, the Arg98 side chain forms hydrogen bonds to residues 308–309. Movement of the Arg98 side chain to interact with DHNA would free the region around residues 305–312 and enable the adoption of a more open conformation for these residues. There is precedence for movement of this region in *Mtb*-MenD; the equivalent region (residues 300-308) has conformational differences between the DHNA-bound and DHNA-free forms. Rearrangement of this part of the allosteric site upon ligand binding is not unprecedented across other ThDP-dependent decarboxylase superfamily enzymes either. In fact, it can be important for the allosteric mechanism; pyruvate decarboxylase binds the positive allosteric regulator, pyruvate, in a similar structural location to the DHNA binding site in MenD, causing significant rearrangement of the allosteric site region and propagation of allosteric signal to the active site (59).

Exploitation of allosteric sites is an emerging strategy for inhibitor development, but it was not known whether compounds targeting the MenD allosteric site would inhibit growth of whole cells *in vivo*. One advantage of allosteric sites is that they tend to bind relatively hydrophobic compounds, which are associated with cell permeability (31). This is also the case for MenD, which binds hydrophilic molecules in its active site and the relatively hydrophobic DHNA in the allosteric site. Our results show that treatment of *Sau* bacteria with exogenous DHNA inhibits their growth. Furthermore, addition of MK-4 to culture media of DHNA-treated cells results in growth rescue, suggesting that growth inhibition of the bacteria is caused by a lack of menaquinone. While we cannot rule out that DHNA may also target other enzymes within the pathway, these data suggest that inhibition of cell growth may be attributable to inhibition of the *Sau*-MenD enzyme. This is supported by recent findings that show addition of DHNA to culture media increased susceptibility of *Sau* to the antibiotic adjuvant cannabidiol, while MK-4 addition had the opposite effect and prevented CBD-mediated growth impairment (60).

This work demonstrates that despite a lack of strict sequence conservation of the DHNA-binding pocket between *Sau* and *Mtb*, feedback inhibition by DHNA is observed in *Sau*-MenD and the *Sau* MK biosynthesis pathway. Our results suggest MenD may be a suitable target for the development of anti-staphylococcal agents.

## Supporting information

Supplemental Data Merged

## Acknowledgments

This work was primarily supported by the Canterbury Medical Research Foundation (TS and JMJ) with additional funding support from a Marsden Fund from the Royal Society Te Apārangi (NATH, WJ, EMMB, GB, JMJ), the Maurice Wilkins Centre (MWC) for Molecular Biodiscovery (EAW, GB, SSD, JMJ, EMMB) and the Biomolecular Interaction Centre (JT, TMA, JMJ). GB is supported by a Sir Charles Hercus Fellowship through the Health Research Council of New Zealand. This research required the use of the MX2 beamline at the Australian Synchrotron, part of Australian Nuclear Science and Technology Organistion, and made use of the Australian Cancer Research Foundation detector. Access to the Australian Synchrotron was supported by the New Zealand Synchrotron Group Ltd. We acknowledge the kind gift of *S. aureus* strains from Dr Monica Gerth, Victoria University, Wellington.

