## Supplemental Data Merged for "Allosteric inhibition of *Staphylococcus aureus* MenD by 1,4-dihydroxy naphthoic acid: A feedback inhibition mechanism of the menaquinone biosynthesis pathway"

**A**

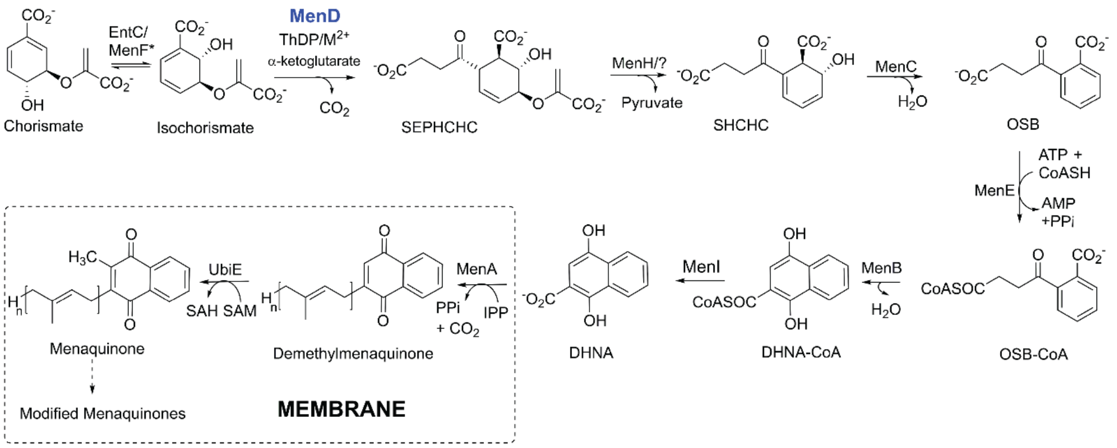


**B**

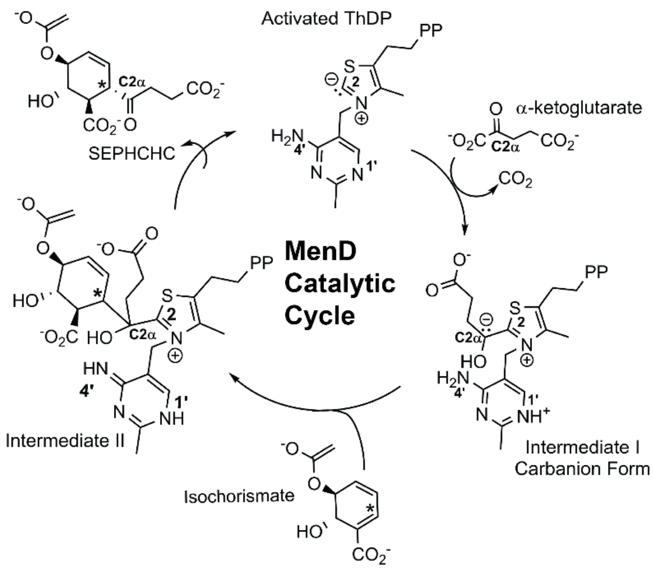


**Figure S1. A.** The classical menaquinone biosynthesis pathway as currently known showing the reactions catalysed by MenA, B, C, D, E, F, H, I and UbiE (sometimes termed MenG) with: SEPHCHC = 2-Succinyl-5-enolpyruvyl-6-hydroxy-3-cyclohexene-1-carboxylic-acid; SHCHC= 2-succinyl-6-hydroxy-2,4-cyclohexadiene-1-carboxylate; OSB= o-Succinylbenzoate; DHNA= 1.4-dihydroxy-2-naphthoic acid; CoA=Coenzyme A; Reactions from MenA onwards are known to be membrane bound. In Mtb DHNA has been shown to inhibit the MenD reaction. **B.** The MenD catalytic cycle in closer detail showing the activated ThDP ylide, the two-step reactions adding each substrate (substrate 1 - α-ketoglutarate/2-oxoglutarate and substrate 2 - isochorismate) and the resulting two ThDP-bound reaction intermediates in the cycle. The aminopyrimidine ring of the ThDP is shown as the AP tautomer in the ThDP ylide, the $APH^+$ tautomer in the Intermediate I carbanion and as the IP tautomer in the Intermediate II form.

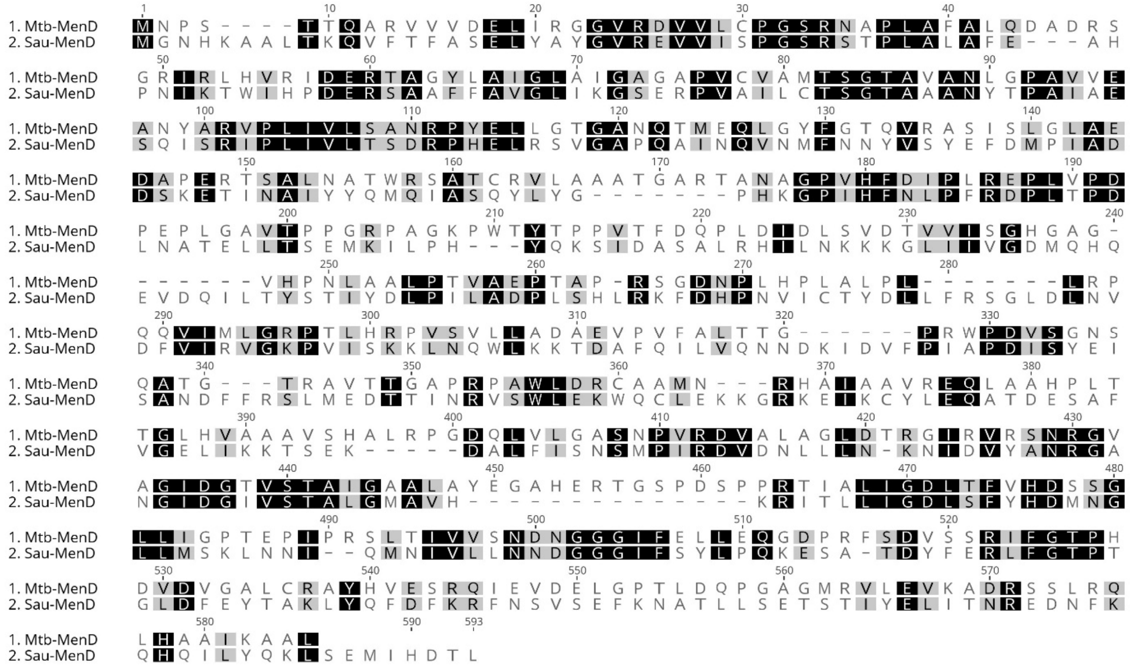


**Figure S2.** MAFFT protein alignment of *Mtb*-MenD and *Sau*-MenD revealed proteins share 28% identity.

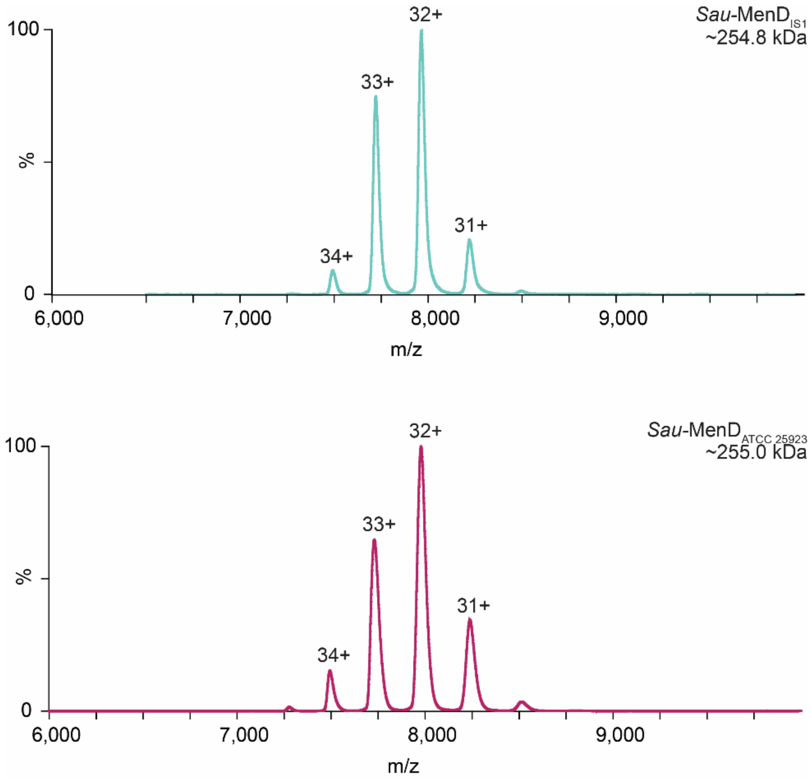


**Figure S3.** Native mass spectra of *Sau*-MenD$_{IS1}$ and *Sau*-MenD $_{ATCC\ 25923}$. The theoretical mass of a tetramer is ~253 kDa for both proteins, with the observed spectra consistent with this oligomeric form. No other species were observed in the mass spectra.

**Table S1.** Primers used in this study**.** All primers were acquired from Integrated DNA Technologies.

| **Primers** | **Sequence (5' - 3')** |
|---|---|
| MenD_Sau_F1 | TTCAGGGCGCCATGGGAAATCATAAAG |
| MenD_Sau_R1 | CGAGTGCGGCCGCAAGCTTATAATGTG |
| menD_Arg98Alafwd | AGCCAAATTAGTGCGATTCCATTAA |
| menD_Arg98Alarev | CGCACTAATTTGGCTTTCAGCAATT |
| menD_Lys283Alafwd | GTTGGGGCGCCAGTGATTTCTAAAA |
| menD_Lys283Alarev | CACTGGCGCCCCAACACGAATTA |
| menD_Lys309Alafwd | AATGATGCGATTGATGTCTTTCCTA |
| menD_Lys309Alarev | ATCAATCGCATCATTGTTTTGCACT |

**Table S2.** *Staphylococcus aureus* and *Pseudomonas aeruginosa* bacteria used in this work.

| **Strain** | **Origin** |
|---|---|
| *Staphylococcus aureus* IS1 | Gift from Dr. Monica Gerth, Victoria University of Wellington |
| *Staphylococcus aureus* ATCC 2923 | ESR Culture Collection New Zealand, NZRM strain number: 917 |
| *Staphylococcus aureus* clinical isolate | Gift from Dr. Monica Gerth, Victoria University of Wellington |
| Methicillin-resistant *Staphylococcus aureus* | ESR Culture Collection New Zealand, NZRM strain number: 1057 |
| *Pseudomonas aeruginosa* clinical isolate | Gift from Dr. Monica Gerth, Victoria University of Wellington |

**Table S3.** *Sau*-MenD$_{IS1}$ data collection and refinement statistics

| | ***Sau*-MenD$_{IS1}$ (7TIN)** |
|---|---|
| **Data collection** | |
| X-ray source | MX2 (λ = 0.95372) |
| Space group | $P2_12_12_1$ |
| Unit cell dimensions (Å) | *a = 81.445, b = 166.489, c = 168.035; α = β = γ = 90.00* |
| Resolution range[a] (Å) | 47.85–2.35 (2.39-2.35) |
| No. of unique reflections | 95884 (4731) |
| Multiplicity | 26.7 (26.9) |
| $R_{merge}$[a] | 0.284 (3.930) |
| $R_{pim}$[a] | 0.056 (0.764) |
| CC ½ [a] | 0.998 (0.346) |
| <I/σ(I)>[a] | 11.5 (1.0) |
| Completeness[a] (%) | 100.0 (99.8) |
| Wilson B (Å$^2$) | 53.09 |
| **Refinement** | |
| Resolution range for refinement [a] | 46.47–2.35 (2.38-2.35) |
| R/$R_{free}$ | 0.1874/0.2355 (0.3013/0.3346) |
| No. of reflections working/test | 95752/4853 (2987/183) |
| No. of non-H atoms | 18089 |
| Protein atoms/waters | 17728/248 |
| Ligands/Ions | 4 TPP; 7 GLY/4 Ca ions; 1 Cl ion; 4 Na ions |
| **B factors** (Å$^2$) | |
| Protein | 56.46 |
| Water | 52.62 |
| Ligands/Ions | 50.46/54.45 |
| Molprobity Score | 1.66 (98th %ile) |
| Estimated coordinate error (Å) | 0.34 (max. likelihood) |
| Clashscore | 3.77 (99th %ile) |
| Bond lengths RMSZ | 0.0029 |
| Bond angles RMSZ | 0.58 |
| **Ramachandran** (%) favoured/allowed/outliers | 97.02/2.98/0 |

[a]Values in parentheses correspond to the highest resolution shell.

Crystallographic data and structure files are available from the Protein databank under the PDB code: 7TIN. The other data from this work is given in the graphs and tables presented in the main manuscript and supplemental information with raw data available on request to the authors.
